# A CTLA-4 nanobody improves the immunity of mice against challenges with *Staphylococcus aureus* and *Streptococcus agalactiae*

**DOI:** 10.1101/2021.03.05.434056

**Authors:** Peng Wu, Ningning Yang, Mingguo Xu, Chuangfu Chen

## Abstract

Cytotoxic T lymphocyte associated antigen 4 (CTLA-4), also known as CD152, is a transmembrane receptor and leukocyte differentiation antigen on T cells that participates in the negative regulation of immune responses. CTLA-4 Ig can effectively and specifically inhibit cellular and humoral immune responses in vivo and in vitro, and is therefore, considered to be a promising new immunosuppressive antibody. In this study, we investigated the role of CTLA-4 nanobody in immunity. We purified recombinant CTLA-4 protein and constructed a phage display nanobody library. After screening the library, we obtained a nanobody with high affinity for the CTLA-4 protein. The nanobody was expressed and purified and the specific high-affinity for CTLA-4 confirmed by enzyme-linked immunosorbent assay. The nanobody was shown to enhance the activity and nitric oxide (NO) production of bone marrow derived dendritic cells (DCs) as well as their ability to capture foot-and-mouth disease virus (FMDV). The CTLA-4 nanobody also improved the immunity of animals after challenge with *Staphylococcus aureus* and *Streptococcus agalactiae*, thus indicating the potential of the CTLA-4 nanobody to improve cellular immunity and enhance immune responses.

## Introduction

CTLA-4, which is a homologue of CD28, is located on the same chromosome band as the CD28 gene [1-3]. Both CD28 and CTLA-4 function as receptors for B7 ligands that are expressed on antigen-presenting cells (APCs) [3]. Despite clear evidence that CTLA-4 is involved in the downregulation of activated T cells, the mechanism is still unclear [4-6]. However, two reasonable possibilities have been proposed. First, CTLA-4 may inhibit the function of T cells by inducing apoptosis of activated T cells [1]. Second, CTLA-4 may inhibit T cell activation by preventing early or late events in T cell activation [1]. CTLA-4 usually exists in the form of a dimer, although the mechanism thorough which the dimer combines remains unclear.

Studies have shown that mouse CTLA-4 dimers are bound by non-covalent bonds [5,7]. The function of CTLA-4 as an immunosuppressive checkpoint has attracted much attention [8-11]. A study of the role CTLA-4 in immune escape of *Schistosoma japonicum* and the underlying mechanism showed that the percentage of CD4+CD25+ regulatory T cells and the number of cytokines in the experimental group injected with an anti-CTLA-4 antibody were significantly higher than those in the control group, suggesting that CTLA-4 is beneficial to the immune escape of *Schistosoma japonicum* [12,13]. As an immunosuppressive receptor on T cells, CTLA-4 participates in the induction of T cell anergy and the inhibition of T cell activation [14-17]. Immunosuppressive receptors are expressed on activated T cells, B cells and macrophages [18].

Antibody blockers of immunosuppressive receptors such as CTLA-4 can promote the transformation of CD8+T cells into memory CD8+T cells, and then rapidly destroy pathogenic microorganisms or produce antibodies when pathogenic microorganisms or vaccines re-enter are encountered, thus demonstrating their capacity to improve immunity in vivo [19,20]. The CTLA-4 nanobody acts directly on immune cells and releases the suppressed state of immune cells. Therefore, in the present study, we generated a CTLA-4 nanobody and confirmed its ability to improve immunity.

## Materials and methods

### Cells and plasmids

HEK293T cells, Madin Darby Bovine Kidney (MDBK) cells, sheep kidney cells, hamster kidney (BHK-21) cells, and FMDV, which was expressed in phages, were preserved by the zoonotic laboratory of Shihezi University (Shihezi, Xinjiang, China). *Staphylococcus aureus* and *Streptococcus agalactiae* strains were isolated and preserved by the zoonotic laboratory of Shihezi University, and cells were cultured in Dulbecco’s modified Eagle’s medium (DMEM; Thermo Fisher Scientific, Massachusetts, USA) supplemented with 10% fetal bovine serum (10% FBS; Thermo Fisher Scientific) at 37°C under 5% CO_2_. *E. coli* strain TG1 was cultured on lysogeny broth (LB) plates overnight at 37°C.

### Ethics statement

Female BALB/c mice (aged 6–8 weeks) were purchased from the Animal Experiment Center of Xinjiang Medical University (Urumqi, China). A healthy young (aged 2 years) male alpaca was obtained from the Key Laboratory of Local and National Diseases High Incidence of Shihezi University. All the experimental procedures involving animals were approved by the Animal Experimental Ethical Committee Form of the First Affiliated Hospital of Medical College, Shihezi University (No. A 280-167).

### Cloning, expression and purification of recombinant CTLA-4

Based on the sequence in the Uniprot database (RefSeq. NM-005214.4), CTLA-4 was synthesized and cloned into the pCDNA3.1+ plasmid (Sangon Biotech, Shanghai, China) with *EcoR* I and *Hind* III restriction enzymes (TaKaRa, Japan) to generate the pCDNA3.1-CTLA-4 plasmid. Positive clones were screened by PCR, and the plasmids were extracted. The plasmid (10 μg) and LipofectamineTM2000 (20 μL; Thermo Fisher Scientific) were added to 1 × 10^6^ concetration of HEK293T cells; the cells were cultured at 37°C under 5% CO_2_ atmosphere. After 72 h, the cells were collected and sonicated for 10 min before centrifugation at 16 000 r/min for 20 min. The lysate supernatant was collected and the protein were separated by sodium dodecyl sulfate polyacrylamide gel (15%) electrophoresis (SDS-PAGE) electrophoresis. CTLA-4 protein was then purified using a protein purification kit (Cwbio, Beijing, China).

### Western blotting analysis of CTLA-4

Expression of recombinant CTLA-4 protein was analyzed by Western blotting. Briefly, the recombinant CTLA-4 protein was purified from the cell lysates and separated by 15% SDS-PAGE. Protein was then transferred to a nitrocellulose (NC) membrane, which was blocked in 5% skimmed milk powder (BD, USA) for 2 h at room temperature (RT). The membrane was washed four times using phosphate-buffered saline-2% Tween 20 (PBST, Solarbio, Beijing, China) and incubated at RT for 2 h with mouse anti-CTLA-4 monoclonal antibody (diluted: 1:3 000; Abcam, Cambridge, UK) as the primary antibody. The membrane was washed and then incubated at RT for 1 h with horseradish-peroxidase (HRP)-conjugated goat anti-mouse (IgG) polyclonal antibody as the secondary antibody (1:5 000; Abcam). After three washes, the immunoreactive band was visualized using a DAB chromogenic reagent kit (Solarbio).

### Alpaca immunization and antibody titer determination

Before immunization, 20 mL of peripheral blood was collected from the alpacas. The serum was separated and frozen at -20°C. On days 28, 49, and 70, the alpacas were immunized subcutaneously and intradermally with 3 mL of 0.5 mg CTLA-4 protein and an equal volume of complete Freund’s adjuvant (0.5 mg; Sigma–Aldrich, USA). On days 91 and 112, the alpaca was immunized subcutaneously and intradermally with 3 mL of 0.25 mg CTLA-4 protein and an equal volume of incomplete Freund’s adjuvant. On day 7 after each immunization, 20 mL peripheral blood was collected from the alpaca. The serum was separated and frozen at -20°C. Anti-CTLA-4 antibody titers were determined by enzyme-linked immunosorbent assay.

### Phage library construction

For phage library construction, the VHH gene was amplified by nested PCR. Total mRNA was extracted from alpaca peripheral blood mononuclear cells (PBMCs; RNApure Tissue&Cell Kit, Cwbio), and transcribed into cDNA (HiFiScript cDNA Synthesis Kit, Cwbio). Using cDNA as the template, the gene fragment was amplified using the following specific primer pair: CAL-F 5’-GTCCTGGCTGCTCTTCTACAAGG-3’ and CAL-R 5’-GGTACGTGCTGTTGAACTGTTCC-3’. The amplified PCR product was purified by 1% agarose gel electrophoresis and used as template for the second PCR with the following primers: VHH-F 5’-GATGTGCAGCTGCAGGAGTCTGGRGGAGG-3’ and VHH-R 5’-CTAGTGCGGCCGCTGAGGAGACGGTGACCTGGGT-3’. The PCR product and the pMECS phagemid vector were digested sequentially with *Pst* I and *Not* I (TaKaRa, Dalian, China) and then ligated by T4 DNA ligase at 16°C overnight. The construct was then transformed into *E*.*coli* TG1 cells and cultured on LB plates containing ampicillin (100 µg/mL). After 37°C incubation overnight, resistant colonies were picked from plates. The size of the library was calculated according to the number of colonies after 10^4^, 10^5^, or 10^6^ dilution. Twenty single clones were randomly picked from the gradient dilution plates of each library, and colony PCR was performed using the following primers: MP57-F 5’-TTATGCTTCCGGCTCGTATG-3’ and MP57-R 5’-CTAGTGCGGCCGCTGAGGAGACGGTGACCTGGGT-3’.

### Screening of Special VHH against CTLA-4

The VHH phage display library was panned in three consecutive rounds. CTLA-4 protein (10, 5 or 1 μg/mL) in carbonate buffer (pH 9.6) was used to coat 96-well plates. Briefly, wells were filled with CTLA-4 solution (100 μL) and incubated overnight at 4°C. After three washes with 1% PBST (300 μL), the plates were blocked with 5% skimmed milk powder (200 μL) for 2 h at 37°C. After washing with PBST, phage library solution was added to each well and incubated for 2 h at 37°C. Wells were then washed 10 times with PBST to remove unbound phages (washing times were increased in the second (15 times) and third (20 times) rounds to reduce non-specific binding). Glycine-HCl (pH 2.2, 100 μL) was added for 10 min to elute target-bound phages and immediately neutralized with 20 μL 2 M Tris-HCl (pH 9.1), followed by incubation at 37°C for 30 min. Panning output was assessed on the following day. Each round of library screening required amplification and titration.

### Construction, expression and purification of the nanobody

To verify the expression level and activity of the nanobody, the CTLA-4 nanobody sequence was cloned into the pET28a-SUMO vector for intracellular expression. After disruption by ultrasonication, the protein was purified using nickel charged columns (this was completed with the assistance of Bard Biotechnology). The CTLA-4 nanobody was analyzed by SDS-PAGE and Western blotting.

### CTLA-4 affinity measurements

Dilute CTLA-4 eukaryotic expression protein was coated onto 96-well (100 ng/well). The plates were blocked overnight at 4°C with 5% skimmed milk powder. CTLA-4 protein was coated onto 96-well-plate (100 ng/well). The plates were blocked overnight at 4°C with 5% skimmed milk powder. The plates were washed three times with PBST before adding the CTLA-4 nanobody (50 µL per well, diluted 1:30, 1:60, 1:120, 1:240, 1:480, 1:920, 1:1 840, and 1:3 680). The plates were incubated for 2 h at 37°C and then washed three times with PBST. The plates were then incubated with 100 µL concentration in each well mouse anti-alpaca monoclonal antibody (1:2000; Thermo Fisher Scientific) for 1 h at 37°C. After three washes with PBST, the plates were incubated with 100 µL/ well HRP-conjugated rabbit anti-mouse antibody (1:10 000; Thermo Fisher Scientific) for 1 h at 37°C. The plates were then washed three times with PBST before adding TMB (100 μL; Solarbio) substrate. The reaction was stopped by the addition of 0.5-2 M H_2_SO_4_ (100 μL; Solarbio). Finally, the OD450nm was measured in each well.

### Cytotoxicity assay

BHK-21 cells, MDBK cells and sheep kidney cells were plated in 96-well plates at 6×10^3^ cells per well. After 3 h, after the cells had adhered, the CTLA-4 nanobody was added at 5 µg/mL, 10 µg/mL, 20 µg/mL or 40 µg/mL. MTS reagent (BioVision; Milpitas, USA) was added (20 µL/well) and incubated at 37°C for 3 h. After shaking, the OD492nm was measured in each well.

### Nanobody and bacteriophage NO detection

Mouse DCs were isolated according to the method described by Inaba et al. [21,22]. Briefly, cells were flushed from the tibias and femurs of BABL/C mice, purified and cultured with 10% FBS containing GM-CSF (10 ng/mL) and IL 4 (10 ng/mL) (all PeproTech, USA). FBS (10%) was added on days 2 and 4 and the cells were collected on day 6. The NO content in DCs was evaluated using a NO detection kit (Solarbio). Briefly, 1 x 10^5^ DCs were added to the 96-well cell culture plates under 5% CO_2_ at 37° incubation for 3 h before the addition of FMDV (1 µg/mL). The cells were then incubated for a further 24 h. The CTLA-4 nanobody was diluted into the wells using 10% FBS culture solution, and the final concentration was adjusted to 10, 20, 40, or 80 µg/mL. The CTLA-4 nanobody dilutions were added to the cells (three replicate wells per sample) and cultured under 5% CO_2_ under 37° for 38 h. In the control group, 100 μL extraction solution, 50 μL reagent one, and 50 μL reagent two (Solarbio) were added. In the experimental group, 100 μL sample, 50 μL of reagent one, and 50 μL reagent two were added. After mixing, the plate was left to stand at RT for 15 min and the OD550nm was measured in each well.

### Cytokine and NO detection

Female BALB/c mice (age: 6-8-week; n = 8 per group) received 0.1 mg (200 μL) of CTLA-4 nanobody or an equal volume of the control group (PBS) via the intramuscular route. Three days after injection, blood was collected and the serum was separated for analysis of the levels of IL-4, IFN-γ and NO (Solarbio).

### *In vivo* challenge experiment

Female BALB/c mice (age: 6-8-week; n = 8 per group) received 0.1 mg (200 μL) of CTLA-4 nanobody or an equal volume of the control group (PBS) via the intramuscular route. Three days after immunization, the mice were challenged with 1.9 × 10^9^ CFU *Staphylococcus aureus* (150 μL) or 5.1 × 10^10^ CFU *Lactococcus agalactiae* (200 μL). After 24 h, the number of surviving mice was observed.

### Statistical analysis

All data were expressed as mean ± standard deviation (SD). Statistical analysis was performed using GraphPad Prism 8 software. Significant differences between groups were evaluated by Mann–Whitney U-test. The threshold for statistical significance was set at P < 0.05.

## Results

### Preparation of CTLA-4 antigens

The CTLA-4 gene was amplified (Fig. 1A) and cloned into the pcDNA3.1 vector (Fig. 1B). The resulting pCDNA3.1-CTLA-4 recombinant plasmid was then transfected into HEK293 for expression of the recombinant CTLA-4 protein. SDS-PAGE analysis revealed that the recombinant protein was of high quality and the purity exceeded 90% (Fig. 1C). The molecular weight of the recombinant CTLA-4 protein was determined to be approximately 23 kDa by Western blot analysis (Fig. 1D).

**Fig. 1.**
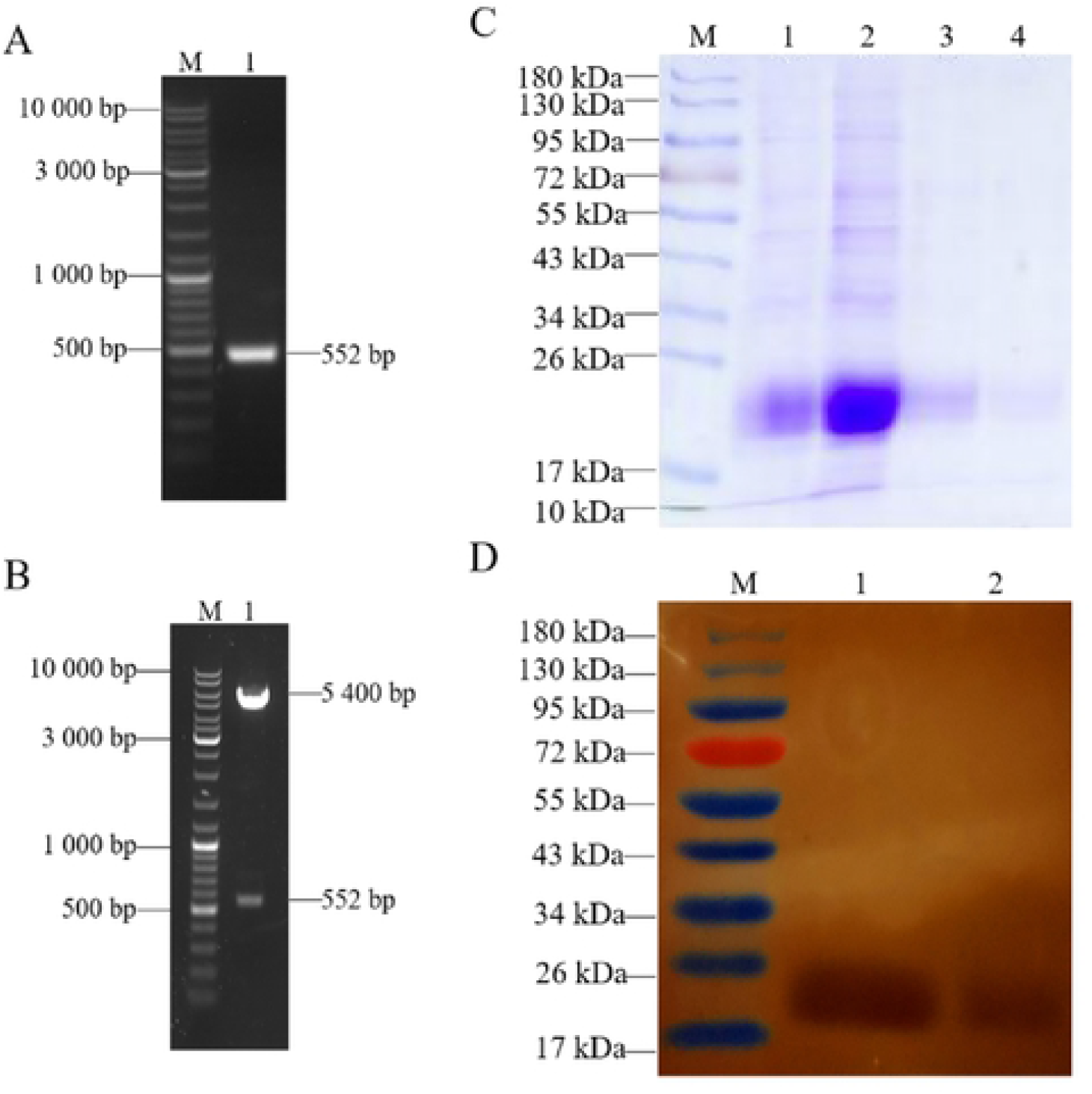
Preparation of CTLA-4 antigens. (A) SDS-PAGE of CTLA-4 gene amplification. M: DNA Marker 10 000; Lane 1: Positive clones. (B) Recombinant pCDNA3.1-CTLA-4 plasmid digested identification with the *EcoR* I and *Hind* III restriction enzymes. M: DNA Marker 10 000; Lane 1: Double enzyme digestion of positive clones. (C) SDS-PAGE results of CTLA-4 protein. M: protein marker (10–180 kDa); Lanes 1–2: Recombinant CTLA-4 protein (23 kDa); Lanes 3–4: Purified CTLA-4 recombinant protein (23 kDa). (D) Western blot of the recombinant CTLA-4 protein. M: protein marker (10–180 kDa); Lanes 1–2: CTLA-4 protein specific band (23 kDa).

### Phage display library construction

The alpaca was immunized six times with CTLA-4 protein. Analysis of the serum revealed substantially elevated antibody titers (Fig. 2A). A gene fragment of approximately 400 bp (Fig. 2B) obtained by PCR was successfully cloned into the pMECS phagemid vector. Phagemids were then transformed into *E. coli* TG1 cells and the size of the library reached 3.2×10^9^ colonies (Fig. 2C). Twenty individual clones were selected randomly for PCR analysis, which showed a library insertion rate of 100% (Fig. 2D). All these results suggested that we had constructed a high-quality immunized phage display library.

**Fig. 2.**
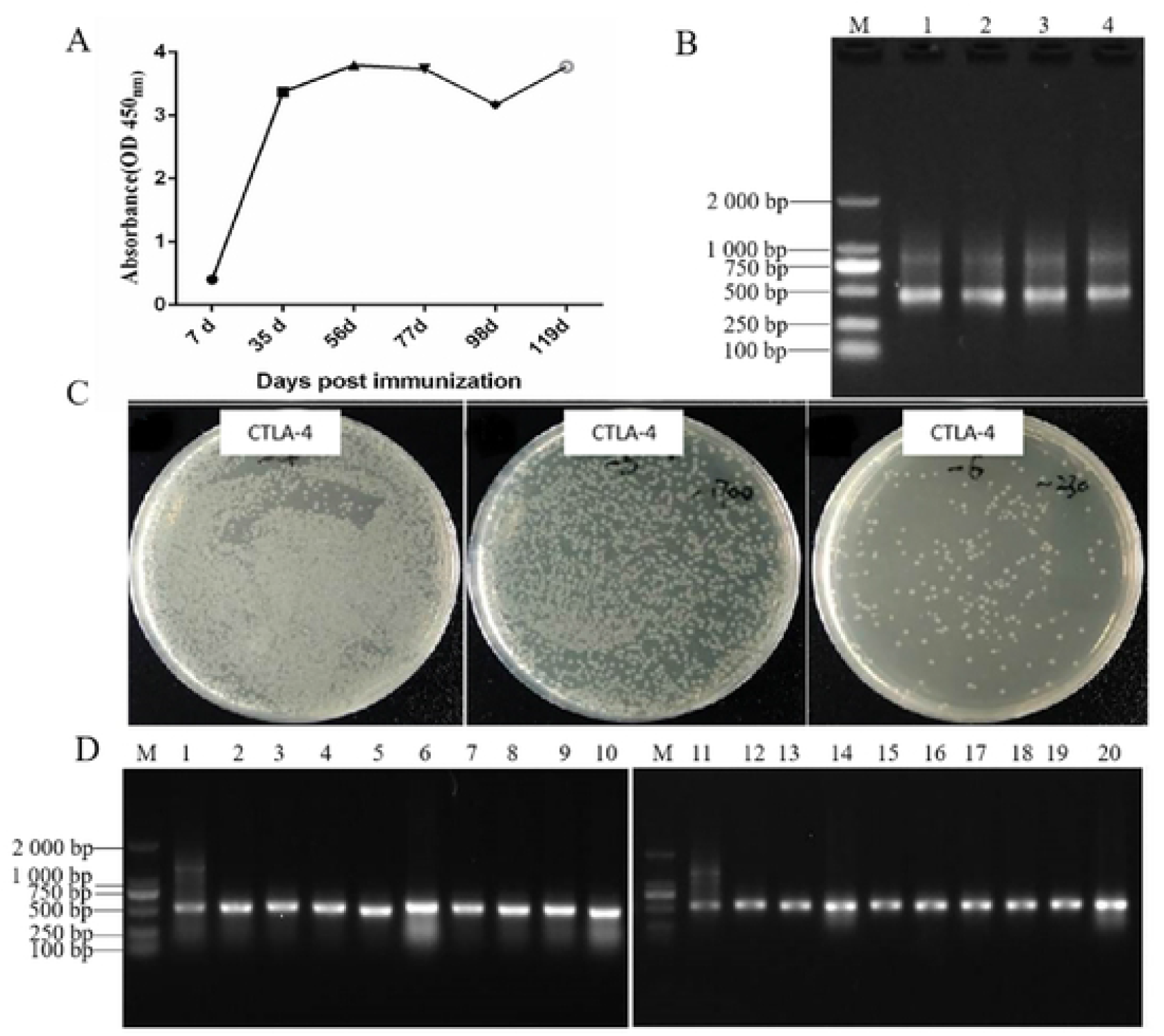
Construction and identification of CTLA-4 nanobody library. (A) CTLA-4 serum titer in the immunized alpaca. (B) Nested PCR amplification of VHH and the 400 bp sequences that were obtained. M: DNA Marker 2000; Lanes 1–4: Positive clones. (C) The library size was measured by counting the number of clones after 10^4^, 10^5^, or 10^6^ dilution. (D) PCR identification Library. M: DNA Marker 1000; Lanes 1–20: Positive clones. Screening of Special VHH Against CTLA-4

As shown in Table 1, the VHH phage display library was panned for three consecutive rounds and the enrichment in each round of bio-panning was calculated according to the following formula: Recovery = output phage (cfu)/ input phage (cfu). cfu: colony-forming unit.

**Table 1.** Selective enrichment of nanobodies from the libraries during panning.

| Round of<br>panning | Input<br>phage<br>(cfu) | Output<br>phage<br>(cfu) | Recovery rate |
| --- | --- | --- | --- |
| Round 1 | $1.0 \times 10^{13}$ | $2.6 \times 10^7$ | $2.6 \times 10^{-5}$ |
| Round 2 | $2.1 \times 10^{12}$ | $1.0 \times 10^8$ | $4.8 \times 10^{-5}$ |
| Round 3 | $1.0 \times 10^{12}$ | $5.8 \times 10^8$ | $5.8 \times 10^{-4}$ |

### Construction, expression, and purification of the nanobody

We successfully constructed and obtained the pET28a-VHHCTLA-4 prokaryotic expression strain. According to SDS-PAGE and Western bot analysis, the molecular weight of the nanobody product was determined to be approximately 23 kDa (Fig. 3).

**Fig. 3.**
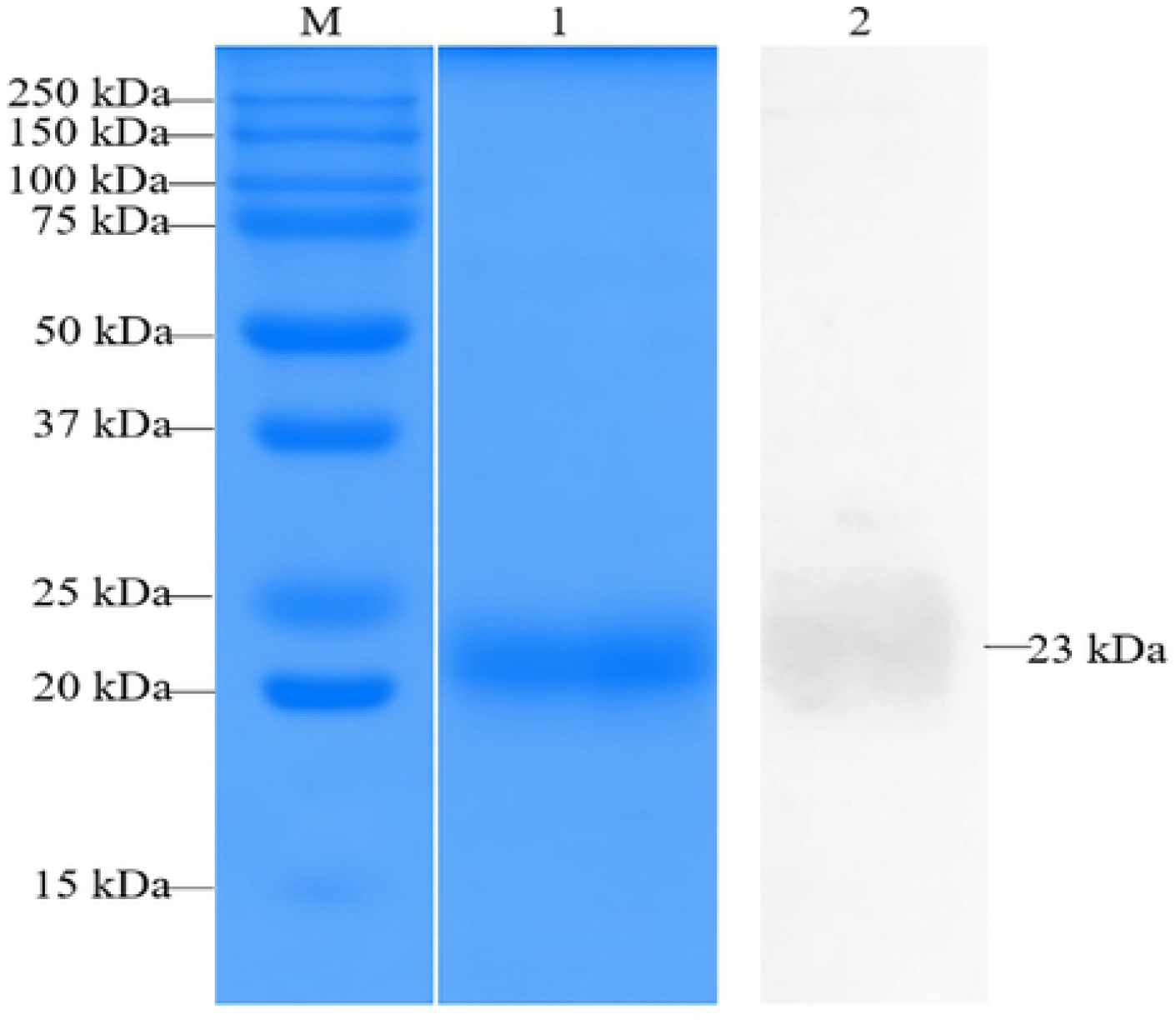
SDS-PAGE and Western blot analysis of the expression and purification of CTLA-4 recombinant protein. M: protein marker (10–250 kDa); Lane 1: SDS-PAGE analysis of the purified recombinant CTLA-4 protein (23 kDa); Lane 2: Western blot analysis of the purified recombinant CTLA-4 protein.

### Affinity and cytotoxicity of the CTLA-4 nanobody

The affinity of the CTLA-4 nanobody for CTLA-4 was assessed by sandwich ELISA. We obtained only one nanobody with high affinity, and the OD450nm was still higher than that of the control group when diluted 3 680 times. This observation indicated a high level of the nanobodies remained bound to the CTLA-4 receptor after dilution 3 680 times (Fig. 4A). In addition, cytotoxicity assays showed that the nanobodies did not mediate cytotoxicity of BHK-21 cells, sheep kidney cells or MDBK cells at any of the concentrations analyzed. These findings confirmed that the CTLA-4 nanobody has no cytotoxic effects on animal cells (Fig. 4B).

**Fig. 4.**
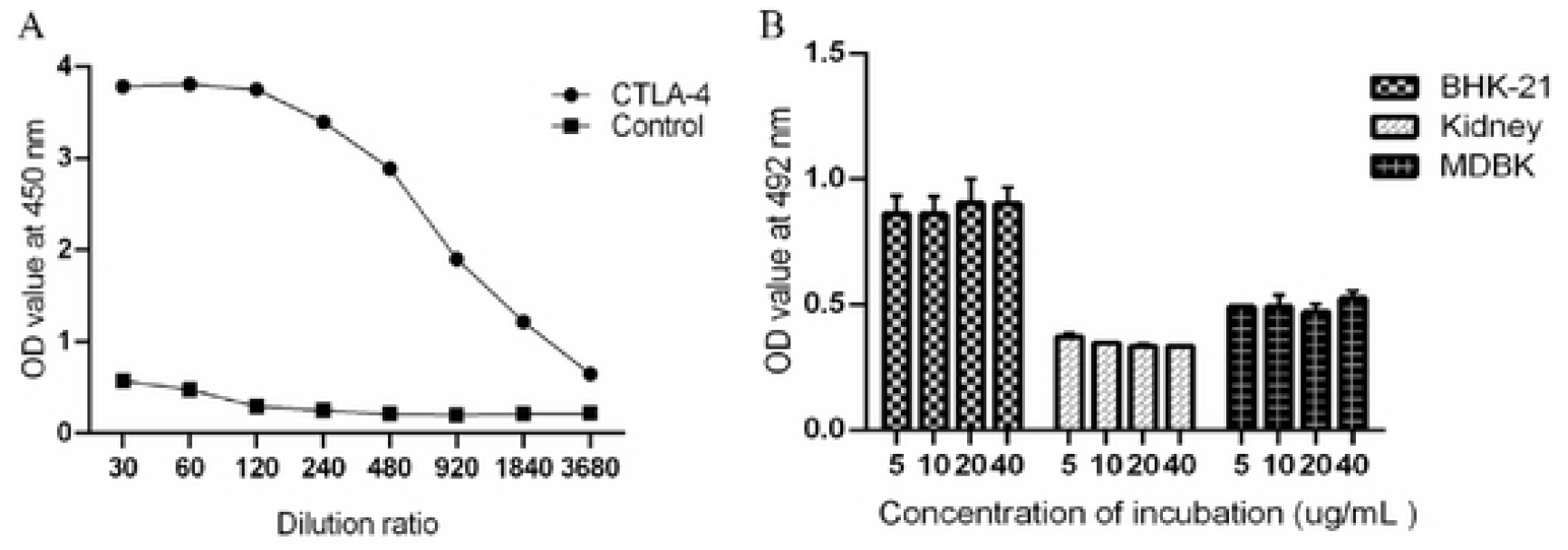
Determination of the affinity and cytotoxicity of the CTLA-4 nanobody. (A) CTLA-4 nanobody affinity. (B) CTLA-4 nanobody cytotoxicity assay.

### CTLA-4 nanobody increased NO production by DCs

Treatment of DCs with the CTLA-4 nanobody induced the production of NO in a concentration-dependent manner (Fig. 5). At 10 µg/mL, the CTLA-4 nanobody had minimal effect on the enhancement of NO production by DCs, while the maximum effect was achieved by treatment with the CTLA-4 nanobody at 80 µg/mL.

**Fig. 5.**
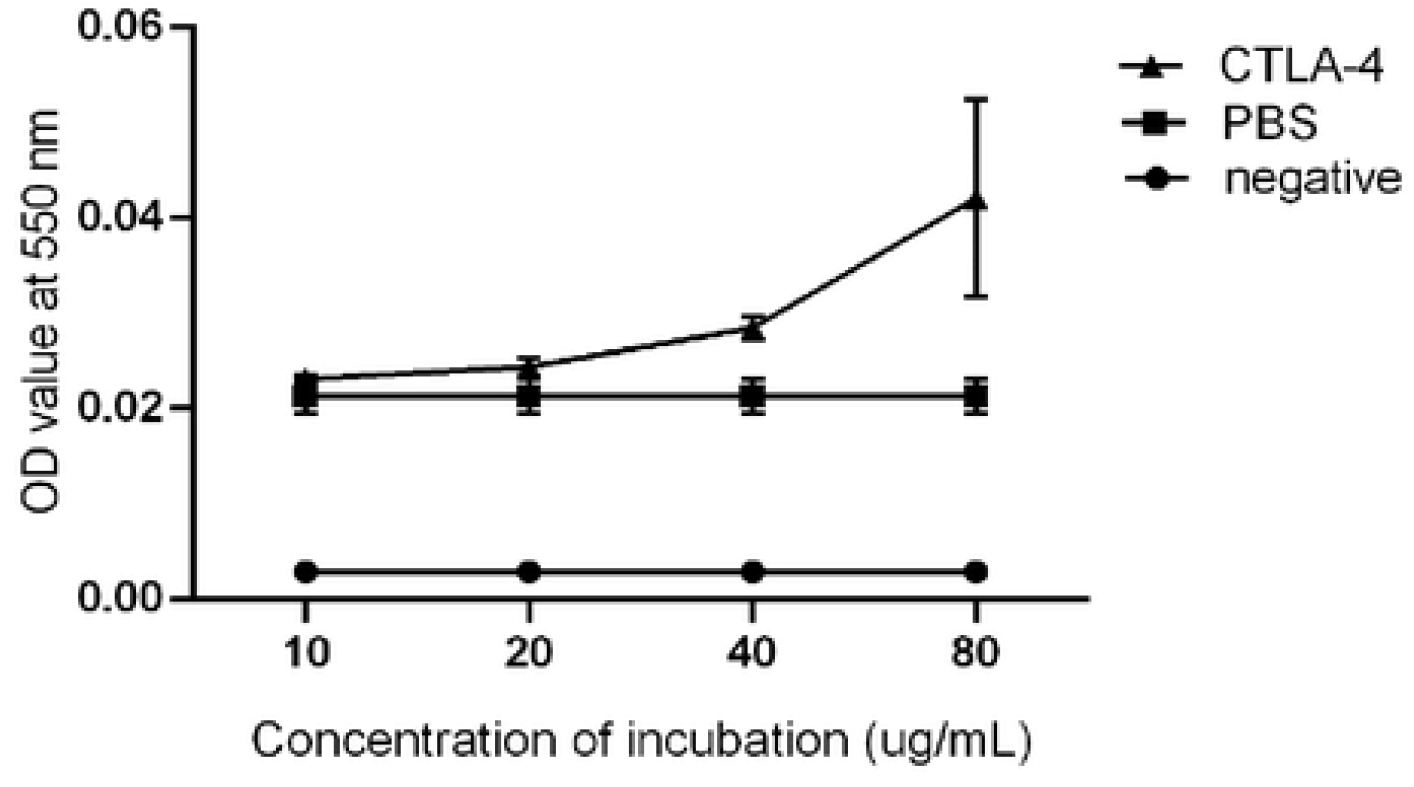
NO determination. Negative: negative control group; PBS: PBS group; CTLA-4: CTLA-4 nanobody group.

### The CTLA-4 nanobody induced a cellular immune response

On the day 3 after the injection of the CTLA-4 nanobody, the levels of cytokine secretion were evaluated by ELISA (n = 8 biologically independent animals per group). The results showed that IFN-γ and NO levels were significantly higher in the CTLA-4 groups than those in the PBS groups (P < 0.001 or P < 0.05), while there was no difference between the groups in IL-4 levels (P < 0.05) (Fig. 6).

**Fig. 6.**
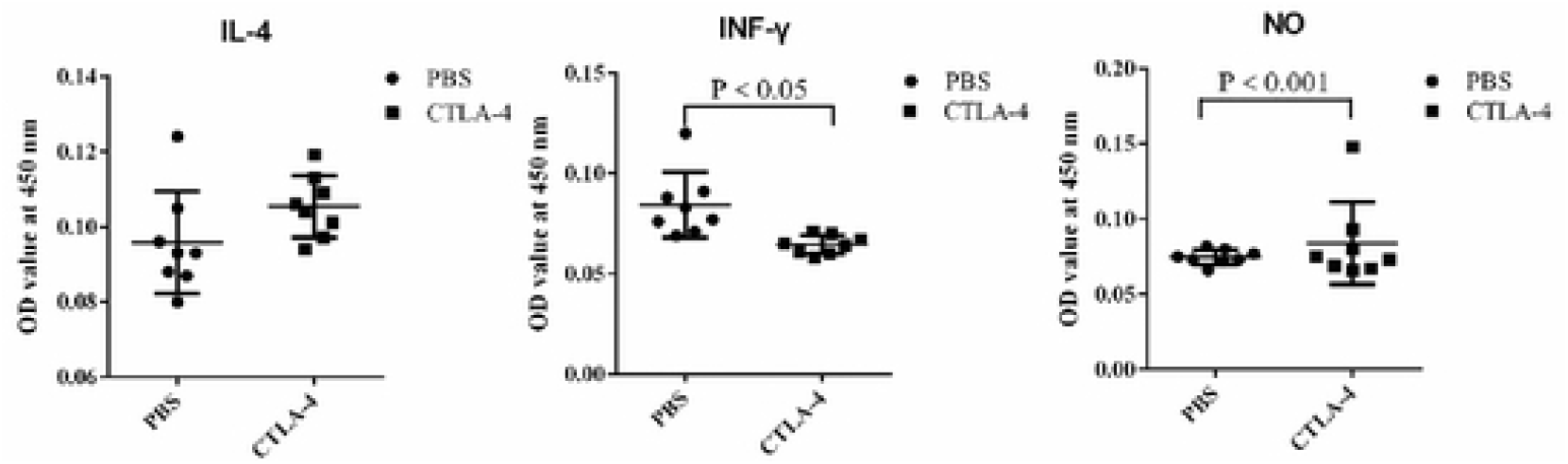
CTLA-4 nanobody induced a cellular immune response. Cellular immune responses were assessed day 3 after the injection with CTLA-4 nanobody of 0.1 mg (200 μL) concetration by ELISA (8 biologically independent animals per group). All experiments were independently performed at least thrice.

### CTLA-4 nanobody improves the survival rate of mice

Administration of the CTLA-4 nanobody improved the partial survival rate of mice, although the effect did not reach the level of statistical significance (Table 2).

**Table 2.**
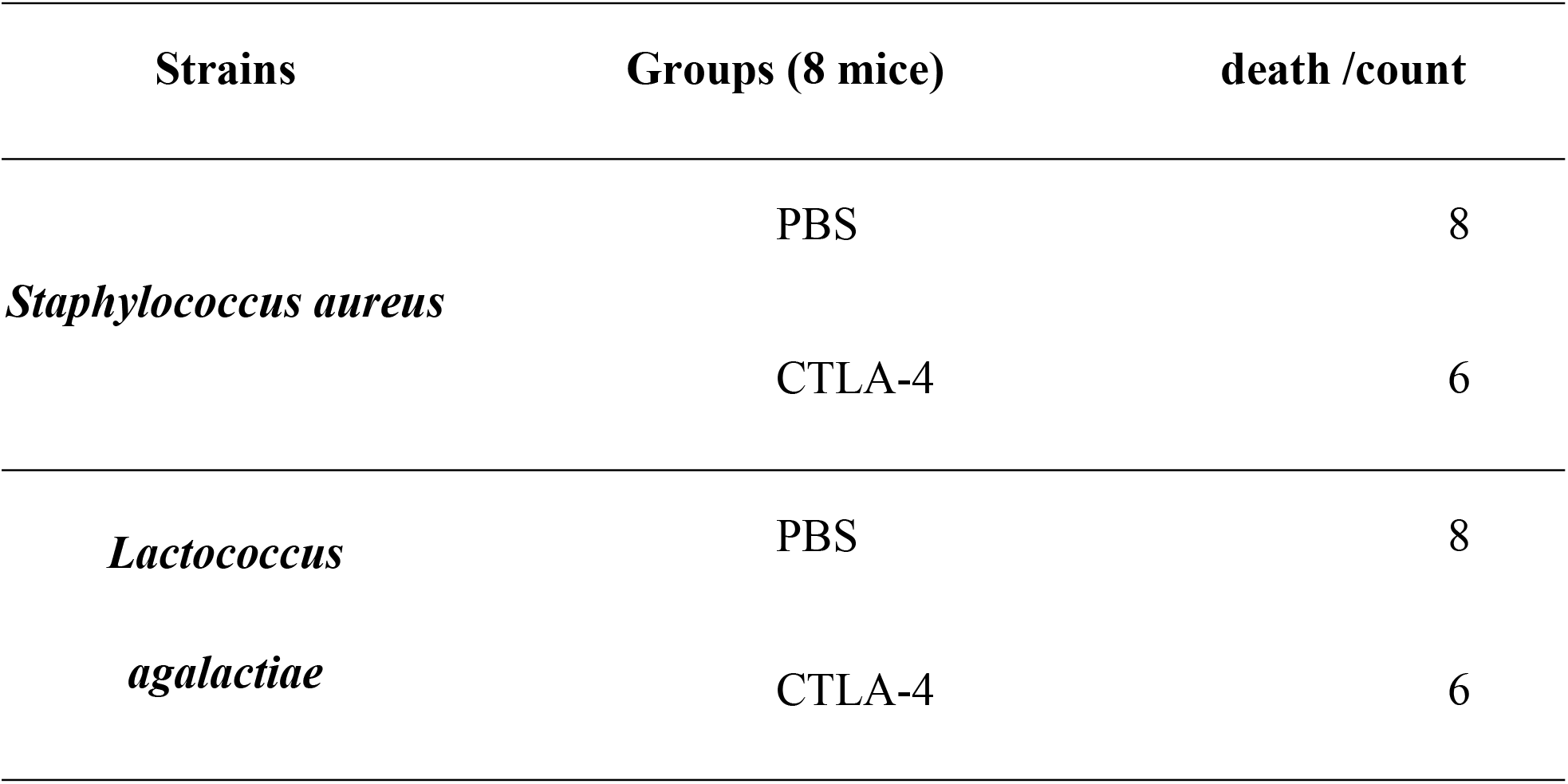
Mice protection study

## Discussion

In this study, we generated and purified a CTLA-4 nanobody that was confirmed to specifically bind to the CTLA-4 protein with a high affinity. Cytotoxicity assays revealed that the CTLA-4 nanobody was non-toxic to cells and was suitable for use in cell experiments. The CTLA-4 nanobody was found to increase the NO production by DCs and enhanced their ability to capture FMDV. In summary, our results indicate the potential of the CTLA-4 nanobody to improve cellular immune responses, with potential protective effects against viral transmission.

The response of animals to antigens in forms such as vaccines is not only related to the quality of vaccines, but also related to the immunity and health status of animals. An immune checkpoint is a receptor that inhibits the activity of immune cells [11]. DCs play a dual role in the regulation of immune responses. Although, in the past, much attention has been centered on their exceptional ability to elicit T cell responses, a crucial role for DCs in immune tolerance has now emerged [23].

When the body is stimulated by bacteria, viruses or other pathogenic microorganisms, these are captured by APCs (especially DCs) and processed for presentation to T cells in the form of an antigen peptide-MHC-II molecular complex. The T cells bind these complexes specifically via their antigen recognition receptors (TCR) [24]. The CTLA-4 inhibitory receptor has been selected for the development of blockers that enhance cellular immune responses. In general, the immune induction period of vaccines is longer than the incubation period of pathogenic microorganisms, thus limiting the protection provided in susceptible animals, and inactivated vaccines cannot prevent reinfection [25]. The use of a CTLA-4 immunopotentiator enhances the response of other immune cells such as suppressed T cells, which indirectly enhances the activity of immune cells, theoretically shortens the induction period of vaccines, and enhances the immune efficacy of inactivated vaccines. Because FMDV is stable in verifying immune cell level, it is often used as a target to verify immune cell activity [26]. Mouse DCs were treated with FMDV followed by incubation with different concentrations of the CTLA-4 nanobody for 24 h. It was found that the NO secretion by DCs reached a maximum following treatment with the CTLA-4 nanobody at 80 μg/mL, thus demonstrating the immune response of DCs was enhanced by the CTLA-4 nanobody.

*Staphylococcus aureus* is the main opportunistic pathogen of humans. It is also one of the most important pathogens causing mastitis in dairy cows [27, 28]. *Streptococcus agalactiae* is one of the most common pathogenic streptococci, and also the most harmful [29,30]. The bacterial strain was first isolated from cows suffering from mastitis. Thus, as highly contagious causes of mastitis in dairy cows, *Staphylococcus aureus* and *Streptococcus agalactiae* are a serious threat to the dairy farming industry [31,32]. In this study, we evaluated the protective effect of the CTLA4 nanobody by challenging mice with wild-type *Staphylococcus aureus* and *Streptococcus agalactiae* isolated from the laboratory. The results of this protection study showed that CTLA-4 nanobody improved the survival rate of mice. Taken together, these findings indicate the CTLA-4 nanobody can be used to promote immune responses to disease-causing pathogens.

## Conclusions

In this study, we generated and purified a CTLA-4 nanobody that was confirmed to specifically bind to CTLA-4 protein with high affinity. Cytotoxicity assays showed that the CTLA-4 nanobody was non-toxic to cells and was suitable for use in cell experiments. The CTLA-4 nanobody was shown to increase NO production by DCs and enhanced their ability to capture FMDV. In summary, our findings indicate the potential of the CTLA-4 nanobody potential to improve cellular immune responses, with the potential to protect against viral transmission.

## Acknowledgment

This research was funded by The Molecular Mechanism of the *Brucella* virulence factors in Persistent infection (Grant no. U1803236); Humanization of 2019 novel coronavirus nanoantibodies (Grant no. ZZZC202084B). The authors would like to thank all the reviewers who participated in the review and MJEditor (www.mjeditor.com) for its linguistic assistance during the preparation of this manuscript.

## Author Contributions

Conceptualization: CC.

Data curation: PW, NY, MX.

Formal analysis: PW. NY.

Funding acquisition: CC.

Investigation: MX.

Methodology: PW, NY, MX.

Project administration: CC.

Resources: CC, PW.

Software: PW, NY.

Supervision: CC, PW.

Validation: CC.

Visualization: MX.

Writing – original draft: PW.

Writing – review & editing: PW, NY, MX.

## References

1. T L Walunas., C Y Bakker., J A Bluestone. CTLA-4 ligation blocks CD28-dependent T cell activation. J Exp Med. 1996, 183, 2541–2550. https://doi.org/10.1084/jem.183.6.2541.

2. P S Linsley, W Brady, M Urnes, L S Grosmaire, N K Damle, J A Ledbetter; CTLA-4 is a second receptor for the B cell activation antigen B7. J Exp Med. 1991, 174, 561–569. https://doi.org/10.1084/jem.174.3.561.

3. Linsley P S., Nadler S G., Bajorath J. et al. Binding stoichiometry of the cytotoxic T lymphocyte-associated molecule-4 (CTLA-4): A disulfide-linked homodimer binds two CD86 molecules. Journal of Biological Chemistry. 1995, 270, 15417–15424. https://doi.org/10.1074/jbc.270.25.15417.

4. Grohmann Ursula., Orabona Ciriana., Fallarino Francesca. et al. CTLA-4-Ig regulates tryptophan catabolism in vivo. Nat Immunol. 2002, 3, 1097–101. https://doi.org/10.1038/ni846.

5. Teft W A., Chau T A., Joaquín Madrenas. Structure-Function analysis of the CTLA-4 interaction with PP2A. Bmc Immunology. 2009, 10, 1–10. https://doi.org/10.1186/1471-2172-10-23.

6. Carreno B M., Bennett F., Chau T A. et al. CTLA-4 (CD152) Can Inhibit T Cell Activation by Two Different Mechanisms Depending on Its Level of Cell Surface Expression. Journal of Immunology. 2000, 165, 1352–6. https://doi.org/10.4049/jimmunol.165.3.1352.

7. Greene J A L., Leytze G M., Emswiler J. et al. Covalent Dimerization of CD28/CTLA-4 and Oligomerization of CD80/CD86 Regulate T Cell Costimulatory Interactions. Journal of Biological Chemistry. 1996, 271, 26762–26771. https://doi.org/10.1074/jbc.271.43.26762.

8. Darlington P J., Kirchhof M G., Criado G. et al. Hierarchical regulation of CTLA-4 dimer-based lattice formation and its biological relevance for T cell inactivation. Journal of Immunology. 2005, 175(2):996–1004. https://doi.org/10.4049/jimmunol.175.2.996.

9. Maurizio Lucchesi., Iacopo Sardi., Gianfranco Puppo. et al. The dawn of “immune-revolution” in children: early experiences with checkpoint inhibitors in childhood malignancies. Cancer Chemotherapy and Pharmacology. 2017. https://doi.org/10.1007/s00280-017-3450-2.

10. Priya S., Xiaofang W., Mousumi B. et al. PD-L1 checkpoint inhibition and anti-CTLA-4 whole tumor cell vaccination counter adaptive immune resistance: A mouse neuroblastoma model that mimics human disease. Plos Medicine. 2018, 15, e1002497. https://doi.org/10.1371/journal.pmed.1002497.

11. Callahan M K., Wolchok J D. Clinical Activity, Toxicity, Biomarkers, and Future Development of CTLA-4 Checkpoint Antagonists. Seminars in Oncology. 2015, 42, 573–586. https://doi.org/10.1053/j.seminoncol.2015.05.008.

12. Hude I, Sasse S., Engert A. et al. The emerging role of immune checkpoint inhibition in malignant lymphoma. Haematologica. 2017, 102, 0–42. https://doi.org/10.3324/haematol.2016.150656.

13. Scheipers P., Reiser H. Role of the CTLA-4 receptor in T cell activation and immunity. Physiologic function of the CTLA-4 receptor. Immunologic Research. 1998, 18,103–115. https://doi.org/10.1007/BF02788753.

14. Romano Audrey., Hou Xunya., Sertorio Mathieu. et al. FOXP3+ Regulatory T Cells in Hepatic Fibrosis and Splenomegaly Caused by Schistosoma japonicum: The Spleen May Be a Major Source of Tregs in Subjects with Splenomegaly. PLoS Negl Trop Dis. 2016. https://doi.org/10.1371/journal.pntd.0004306.

15. Tang Chun-Lian., Yang Jin-Feng., Pan Qun. et al. Anti-CTLA-4 monoclonal antibody improves efficacy of the glyceraldehyde-3-phosphate dehydrogenase protein vaccine against Schistosoma japonicum in mice. Parasitol. Res. 2019, 118, 2287–2293. https://doi.org/10.1007/s00436-019-06363-1.

16. Kim K D., Choi J M., Chae W J. et al. Synergistic inhibition of T-cell activation by a cell-permeable ZAP-70 mutant and CTLA-4. Biochemical & Biophysical Research Communications. 2009, 381,355–360.https://doi.org/10.1016/j.bbrc.2009.02.046.

17. Marjaneh Razmara., Brendan Hilliard., Azadeh K. Ziarani. et al. CTLA-4·Ig converts naive CD4+CD25-T cells into CD4+CD25+ regulatory T cells. International Immunology. 2008, 20, 471–483. https://doi.org/10.1093/intimm/dxn007.

18. Valk Elke., Leung Rufina., Kang Hyun. et al. T cell receptor-interacting molecule acts as a chaperone to modulate surface expression of the CTLA-4 coreceptor. Immunity. 2006, 25, 807–21. https://doi.org/10.1016/j.immuni.2006.08.024.

19. Vaughan A N., Malde P., Rogers N J. et al. Porcine CTLA4-Ig lacks a MYPPPY motif, binds inefficiently to human B7 and specifically suppresses human CD4+ T cell responses costimulated by pig but not human B7. Journal of Immunology. 2000, 165, 3175. https://doi.org/10.4049/jimmunol.165.6.3175.

20. Morris Anna B., Adams Layne E., Ford Mandy L. Influence of T Cell Coinhibitory Molecules on CD8 Recall Responses. Front Immunol. 2018, 9, 1810. https://doi.org/10.3389/fimmu.2018.01810.

21. Alice O., Kamphorst. et al. Beyond adjuvants: Immunomodulation strategies to enhance T cell immunity. Vaccine. 2015. https://doi.org/10.1016/j.vaccine.2014.12.082.

22. Inaba K., Inaba M., Romani N. et al. Generation of large numbers of dendritic cells from mouse bone marrow cultures supplemented with granulocyte/macrophage colony-stimulating factor. Journal of Experimental Medicine. 1992, 176, 1693–1702. https://doi.org/10.1084/jem.176.6.1693.

23. Wang W., Li J., Wu K. et al. Culture and Identification of Mouse Bone Marrow-Derived Dendritic Cells and Their Capability to Induce T Lymphocyte Proliferation. Medical science monitor: international medical journal of experimental and clinical research. 2016, 22, 244–250. https://doi.org/10.12659/MSM.896951.

24. He X., Smeets R L., Koenen H J P M. et al. Mycophenolic acid-mediated suppression of human CD4+ T cells: more than mere guanine nucleotide deprivation. Am. J. Transplant. 2011, 11, 439–49. https://doi.org/10.1111/j.1600-6143.2010.03413.x.

25. Sobrino F., Sáiz M., Jiménez-clavero MA. et al. Foot-and-mouth Disease Virus: a Long Known Virus, but a Current Threat. Veterinary Research. 2001, 32, 1–30. https://doi.org/10.1051/vetres:2001106.

26. Teng Z., Sun S., Chen H., et al. Golden-star nanoparticles as adjuvant effectively promotes immune response to foot-and-mouth disease virus-like particles vaccine. Vaccine. 2018, 36. https://doi.org/10.1016/j.vaccine.2018.09.030.

27. Peton V., Le Loir Y. Staphylococcus aureus in veterinary medicine. Infection Genetics & Evolution. 2014, 21, 602–615. https://doi.org/10.1016/j.meegid.2013.08.011.

28. Xihong Z., Zhixue Y., Zhenbo X. Study the Features of 57 Confirmed CRISPR Loci in 38 Strains of Staphylococcus Aureus. Frontiers in Microbiology. 2018, 9, 1591-. https://doi.org/10.3389/fmicb.2018.01591.

29. Long B., Yong-Qing H., Li-Xia F. et al. Isolation and Identification of Streptococcus agalactiae from Mastitis-affected Cows. Animal Husbandry and Feed ence. 2010. https://doi.org/10.16003/j.cnki.issn1672-5190.2010.01.141.

30. Vincent, P.Richards. et al. Comparative genomics and the role of lateral gene transfer in the evolution of bovine adapted Streptococcus agalactiae. Infection Genetics & Evolution. 2011. https://doi.org/10.1016/j.meegid.2011.04.019.

31. Keefe, Greg. Update on control of Staphylococcus aureus and Streptococcus agalactiae for management of mastitis. Vet Clin North Am Food Anim Pract. 2012, 28, 203–216. https://doi.org/10.1016/j.cvfa.2012.03.010.

32. Nunes E L C., Barbosa E V., Folly E. et al. Bovine mastitis: a brief reminder about a potential target for exploring medicinal plants use. Intl Jrl of Medical Plans and Alternative Medicine. 2013, 1, c80–86.

